# Foot–Ground Force Quantifies Impaired Balance Control Mechanisms Post-Stroke

**DOI:** 10.1101/2025.10.05.680543

**Authors:** Kaymie Shiozawa, Rika Sugimoto-Dimitrova, Kreg G. Gruben, Neville Hogan

**Affiliations:** Department of Mechanical Engineering, Massachusetts Institute of Technology, 77 Massachusets Ave, Cambridge, MA, 02139, USA; Department of Mechanical Engineering, University of Wisconsin–Madison, Madison, 1513 University Ave, Madison, WI, 53706, USA; Department of Kinesiology, University of Wisconsin–Madison, Madison, 1300 University Ave, Madison, WI, 53706, USA; Department of Brain and Cognitive Sciences, Massachusetts Institute of Technology, 77 Massachusets Ave, Cambridge, MA, 02139, USA

**Keywords:** stroke, balance, ground reaction force, inverted pendulum, neural control

## Abstract

Approximately 50% of stroke survivors experience lasting balance impairments that persist and that are often managed through compensatory, but suboptimal, strategies. Identifying neuromechanical control changes after stroke could enable more targeted and effective rehabilitation strategies. Computational modeling has begun to uncover balance control strategies in unimpaired adults, but efforts have been limited post-stroke. Here we show one of the first instances of a model of quiet stance that reveals distinct control strategies in post-stroke individuals compared to similarly-aged unimpaired participants. Quiet standing was modeled using a double-inverted pendulum with full-state feedback control. The controller parameters were fit to foot-ground force data collected from 12 post-stroke and 22 similarly-aged unimpaired participants. The best-fit models revealed a joint-torque-coordination pattern in the paretic limb of post-stroke participants that differed substantially from that of the unimpaired participants. The post-stroke participants’ non-paretic limb also showed increased reliance on neural feedback, which may quantify compensatory effort for the altered coordination in the paretic limb. The results demonstrate that model-based analysis of foot-ground force behavior can reveal clinically meaningful insights that are not captured by traditional assessments.

## 1 Introduction

Stroke, or cerebral vascular accident, can disrupt muscle strength, motor control, proprioception, sensory ability, and cognition, leading to a decrease in overall mobility and independence. Approximately 50% of stroke survivors experience lasting disability [1], with postural balance impairments being one of the most well-documented residual deficits [2]. For example, stroke can cause loss of postural stability and increased sway compared to similarly-aged unimpaired controls [3]. Understanding these impairments is critical for developing targeted clinical and rehabilitative interventions to improve balance ability and reduce fall risk in stroke survivors.

Standing can be broadly categorized into unperturbed (quiet) and perturbed standing. Quiet standing is an inherently unstable task that involves controlling body posture to be near upright (i.e. balancing) despite intrinsic sensory and motor noise that promotes toppling. In contrast, perturbed standing refers to the recovery of upright posture in the face of unexpected external disturbances, such as motion of the support surface, foot slipping, or being pushed. While both types of standing are important for understanding postural control and fall risk, this work focuses on quiet standing for several reasons. First, large perturbations induced during experimentation can pose risks to participants with impairments, such as stroke, thus limiting the extent to which real-world disturbances can be safely simulated in a lab. Second, postural responses to perturbations observed in controlled experimental settings may not fully capture natural quiet standing behavior. Third, repeated exposure to external perturbations may evoke adaptation, resulting in behavior different from natural quiet standing.

Many stroke survivors regain the ability to quietly stand unsupported within the first few days after experiencing a stroke [4]. However, despite improvements in functional balance and mobility, certain impairments often persist, such as left-right vertical foot-force mismatch (i.e., weight-bearing asymmetry), reduced activation of key muscles (e.g., paretic-side hamstrings), and increased sway [4–6]. Individuals likely adopt compensatory strategies that allow them to achieve functional balance despite underlying impairments [7]. Nevertheless, approximately 50% of individuals with severe stroke (total anterior circulation infarction) do not regain independent mobility even after more than a month of rehabilitation [4], suggesting the importance of addressing impairments rather than opting for compensation.

While there is a significant body of research on post-stroke balance, much of the existing literature focuses on describing impairments and disabilities rather than uncovering the underlying mechanisms of balance deficit [1, 8–10]. A deeper mechanistic understanding could help inform therapeutic interventions aimed at restoring balance control rather than merely promoting compensatory strategies [7]. One promising approach to address this gap is computational modeling, which has been employed to explore balance control strategies in unimpaired populations [11]. A similar approach could be applied to post-stroke populations to identify control strategies that may be contributing to balance impairment. However, modeling efforts in post-stroke populations remain limited.

Previous work on post-stroke quiet balance modeling includes a musculoskeletal model that characterized postural sway in stroke survivors [12]. The study found that individuals post stroke exhibited altered knee extension control, likely related to compensatory behaviors such as knee hyperextension and forward leaning. However, this model required fitting 45 parameters and only considered center-of-mass (CoM) range and speed, making informative interpretation challenging. Other studies of post-stroke balance have involved perturbations, such as an inverted pendulum model describing balance after transitioning from sitting to standing [13] and work by van Asseldonk and colleagues quantifying the contributions of the paretic and non-paretic limbs to balance control [14].

Traditionally, balance models have often relied on center-of-pressure (CoP) and CoM trajectory analyses. On the other hand, alternative measures of balance may facilitate investigation of post-stroke balance control. Recent research suggests that incorporating the orientation of the foot-ground interaction force—which reflects relative muscle control and is suspected to be disrupted following stroke [7, 15, 16]—may provide insights into the control mechanisms underlying post-stroke balance impairments [17, 18]. The intersection point of the sagittal-plane lines-of-action of foot-ground interaction forces captures the relationship between the CoP position and foot-ground force orientation.

Much like the metacenter of a ship, which must remain above the ship’s center-of-gravity for it to stay upright, the intersection point must always be above the CoM height to stabilize a single inverted pendulum [18]. However, the intersection-point in quiet standing humans has been demonstrated to also move below the CoM height depending on the frequency [17, 19–22]. This behavior could be reproduced by a double-inverted pendulum model, the next simplest model after a single-inverted pendulum [18]. Moreover, the variation of the intersection point’s height across frequency is informative because it is related to the closed-loop standing-balance dynamics via the system frequency response function [23]. Numerical analyses based on a double-inverted-pendulum model stabilized by a full-state feedback controller showed that the distinct patterns of the intersection-point height observed across various balance challenges [17, 19, 20] and different populations [21], including post-stroke individuals [22], were consistent with diverse control strategies. In other words, this method enabled quantification of different joint-torque level control strategies across conditions and ages. Given these findings, the intersection point measure combined with modeling offers a promising approach to quantify post-stroke control of quiet standing.

In this work, we investigated how balance control strategies differed (i) between the non-paretic and paretic limbs in participants with history of stroke and (ii) between similarly-aged individuals with and without history of stroke. We hypothesized that a simple model that can replicate the experimentally observed intersection-point-height data can quantify control strategy differences. Given the significant impairments observed in the paretic limb [1, 8, 10] and the distinct inter-leg differences in the intersection-point height’s frequency dependence reported by Bartloff and colleagues [22], we expected to identify unique control strategies in stroke survivors compared to the similarly-aged participants. Specifically, we anticipated that the paretic limb would exhibit altered control due to neurological damage, while the non-paretic limb may display compensatory adaptations.

To evaluate these hypotheses, published experimental data from 12 post-stroke individuals and 22 similarly-aged unimpaired participants were examined [22, 24]. Each participant was tasked to stand quietly, placing each foot on a force plate. The sagittal-plane components of the foot-ground interaction force were analyzed to assess the frequency-dependent intersection-point height. A double-inverted-pendulum model, incorporating torque-actuated ankle and hip joints corrupted by white noise, was employed to simulate quiet stance, and results were compared to human experimental data. Control parameters yielding the best fit for each participant group’s data were determined, and corresponding stiffness gain matrices were derived from the best-fit parameters.

Model-based analysis of the intersection-point data revealed altered ankle-hip joint coupling in the paretic limb compared to the non-paretic limb and the unimpaired participants. Moreover, both the paretic and non-paretic legs exhibited greater reliance on neural feedback, revealing impairment and compensatory mechanisms at the neural level. These findings underscore the potential of modeling the intersection-point height behavior to uncover underlying impairments that may contribute to quiet-standing balance dysfunction after stroke.

## 2 Results

### 2.1 Intersection Point Height

The experimental quiet-standing data used in this study were obtained from previously published studies [22] and [24]. The experiments by [22] involved 12 participants (4 female, age: 62 *±* 14 years, mean *±* standard deviation) over 6 months post-stroke with chronic hemiplegia who could stand unassisted for at least five minutes with no pain. Control data from 22 unimpaired participants of similar age (11 female, age: 69 *±* 11 years) collected by [24] were used for comparison. The frequency-dependent intersection-point height normalized by center-of-mass (CoM) height with respect to lab frame was computed from the experimental data of center of pressure (CoP) and orientation of the foot-ground force (*θ*_*F*_) following the procedure briefly outlined in Section 4.1.1 and reported in detail in [25].

As previously reported [22], the mean intersection-point heights were above the CoM (i.e. above 1) at lower frequencies and below the CoM height (i.e. below 1) at higher frequencies for unimpaired group and the non-paretic leg, while being distinct and overall highest for the post-stroke non-paretic limb followed by the unimpaired, and then the post-stroke paretic limb (Figure 1).

**Fig. 1:**
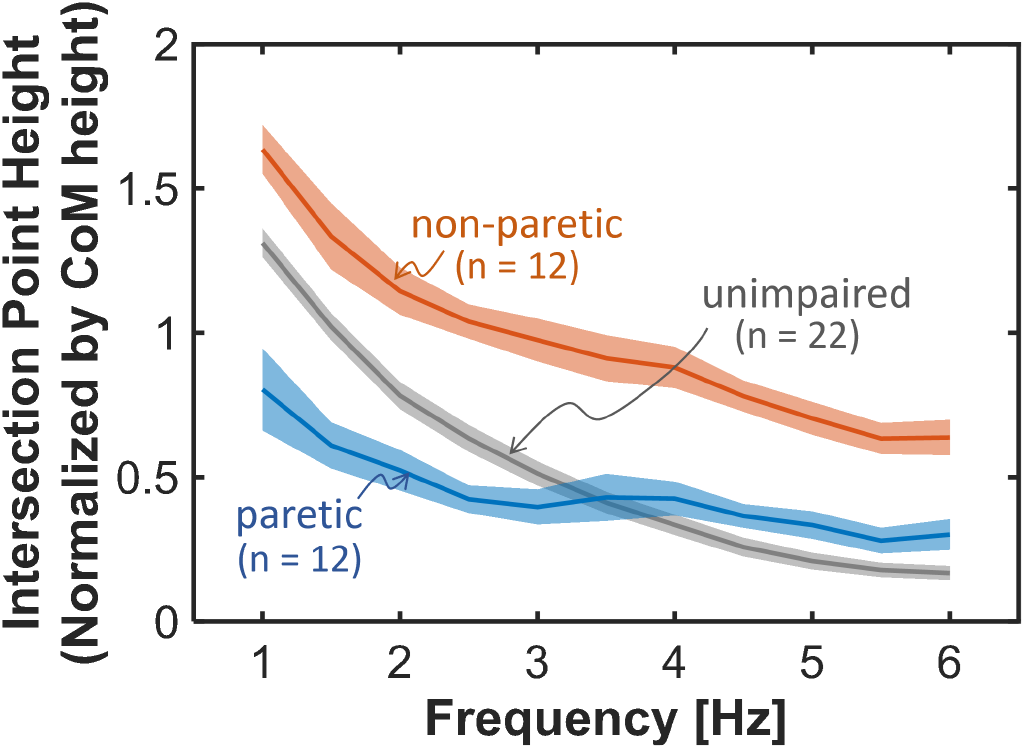
Normalized intersection point height of unimpaired participants (gray), paretic limb of post-stroke participants (blue), and non-paretic limb of post-stroke participants (red). The lines indicate the mean, and the shaded area corresponds to the standard error of the mean, for the 22 unimpaired participants and 12 post-stroke participants.

### 2.2 Best-Fit Control Parameters

The modeling approach used in this work followed a similar procedure to that of previous studies [18, 19, 21]. Because in previous work, it was shown that the single-inverted pendulum could not describe the intersection-point height’s frequency-dependent pattern observed in human participants [18], the next simplest model, the double-inverted pendulum, was used in this study (Figure 2). The model consisted of sagittal-plane ankle and hip joint torques (*τ*_*a*_, *τ*_*h*_) as the input *τ* = [*τ*_*a*_, *τ*_*h*_]^*T*^. The ankle and hip joint angles (*q*_*a*_, *q*_*h*_) and angular velocities 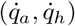 were chosen as the state variables 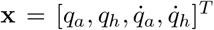. Further details of the model and how the equations of motion, center-of-pressure position, and foot-ground-force orientation were computed can be found in Section 4.2.1 as well as in [18].

**Fig. 2:**
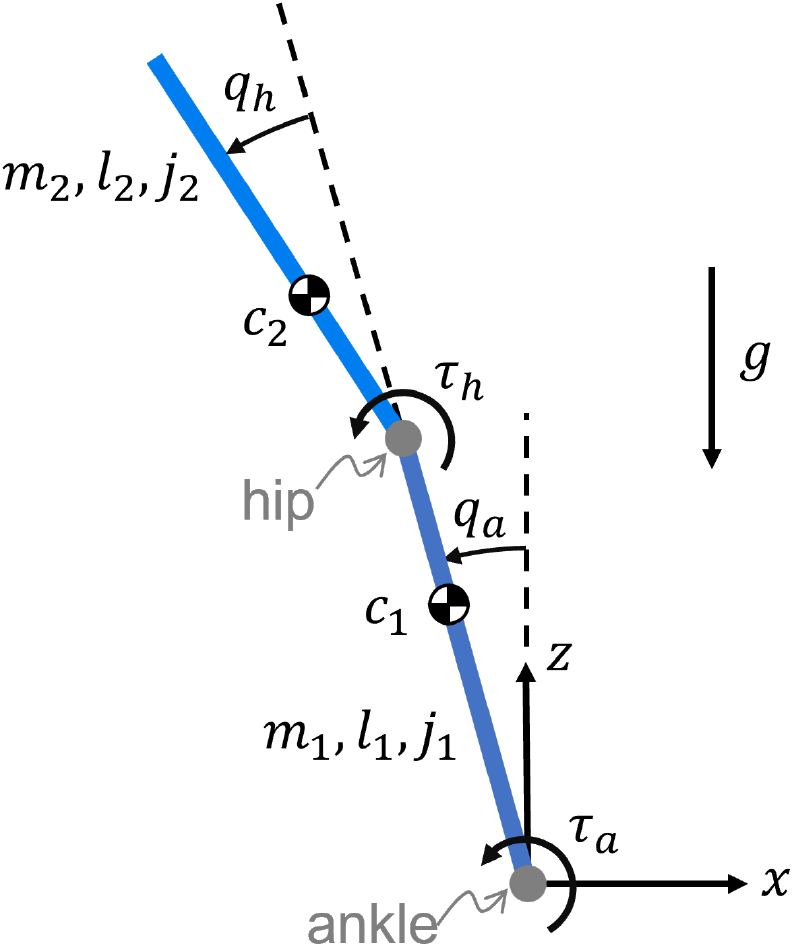
Double-inverted-pendulum model of sagittal-plane standing balance. The two joints represent the ankle and hip, with joint angles *q*_*a*_, *q*_*h*_, and joint torques *τ*_*a*_, *τ*_*h*_, respectively. *τ*_*a*_ is the torque acting on the lower body from the ground, while *τ*_*h*_ is the torque acting on the upper body from the lower body. Link 1 (lower link) represents the legs, with total mass *m*_1_, length *l*_1_, distance from the ankle joint to its CoM *c*_1_, and mass moment of inertia about its CoM *j*_1_. Link 2 (upper link) represents the upper body including the torso, arms, and head, with total mass *m*_2_, length *l*_2_, distance from the hip joint to its CoM *c*_2_, and mass moment of inertia about its CoM *j*_2_. The values used for the lumped model parameters are given in Table 1 in Section 4.2.1.

The controller-specified ankle and hip joint torques were modulated by white, mutually uncorrelated, zero-mean normally distributed noise processes, with standard deviations *σ*_*ankle*_ and *σ*_*hip*_, respectively. The relative strength of the two apparent motor noise processes was described by the noise ratio *σ*_*r*_ = *σ*_*ankle*_*/σ*_*hip*_, and was one of the free parameters that were varied to fit the human intersection-point-height data.

**Table 1.**
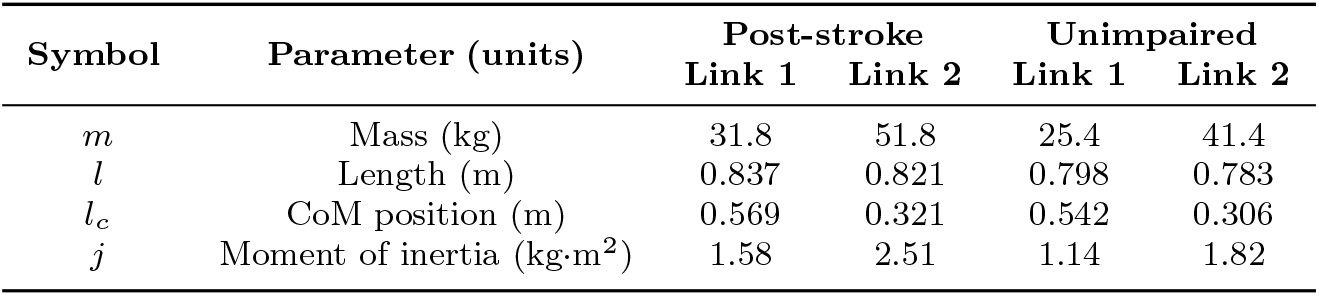
Lumped model parameters.

The apparent full-state-feedback control gains for the best-fit models were identified using the linear-quadratic-regulator (LQR) framework: the double-inverted-pendulum equations of motion were linearized about the upright pose and the control gain matrix **K**_*LQR*_ *∈* ℝ ^2*×*4^ was computed to minimize the cost (*J*) given by:

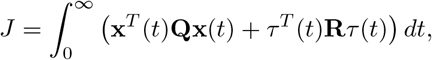

where *τ* = *−***K**_*LQR*_**x, Q** *∈* ℝ^4*×*4^ is the state-cost weighting matrix, and **R** *∈*ℝ^2*×*2^ is the control-input-cost weighting matrix. Inspired by previous work [18, 19, 21], **Q** was parameterized as the identity matrix (*I*_4_) and **R** was parametrized as:

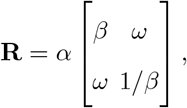

where *α* was fixed as *α* = 10^6^, while *β* and *ω* were varied to find the best-fit model. *β* represents the relative cost on the use of ankle torque versus the use of hip torque (*β >* 1 corresponds to more ankle-torque cost), while *ω* represents the cost on inter-joint coupling (*ω <* 0 encourages positive coupling between ankle and hip torques, while *ω >* 0 encourages negative coupling). Further justification for the chosen parameterization can be found in Section 4.2.2.

A global parameter search was conducted over the parameters *β, ω*, and *σ*_*r*_ to identify the parameter set that best reproduced the experimental mean normalized intersection-point-height curves for each condition (unimpaired, paretic, non-paretic). For each combination of parameters, the intersection-point-height curve was estimated using an analytical method [25]. The parameter combination that yielded the intersection-point-height curve with the lowest root-mean-squared error (RMSE) with respect to the experimental curve was recorded as the best-fitting model for that experimental condition.

The best-fit models’ intersection-point-height curves closely matched the experimental intersection-point-height data (Figure 3). The parameter search yielded the following best-fit parameter sets: for the unimpaired participants, *β* = 0.4, *ω* = *−*0.9, *σ*_*r*_ = 0.6, which yielded *RMSE* = 0.0501; for the paretic limb of the post-stroke participants, *β* = 1.6, *ω* = 0.9, *σ*_*r*_ = 0.9, with *RMSE* = 0.0311; for the non-paretic limb of the post-stroke participants, *β* = 0.7, *ω* = *−*0.9, *σ*_*r*_ = 2.6, with *RMSE* = 0.0571. These best-fit models for the three conditions were compared in subsequent analyses to describe the differences in the apparent balance controllers.

**Fig. 3:**
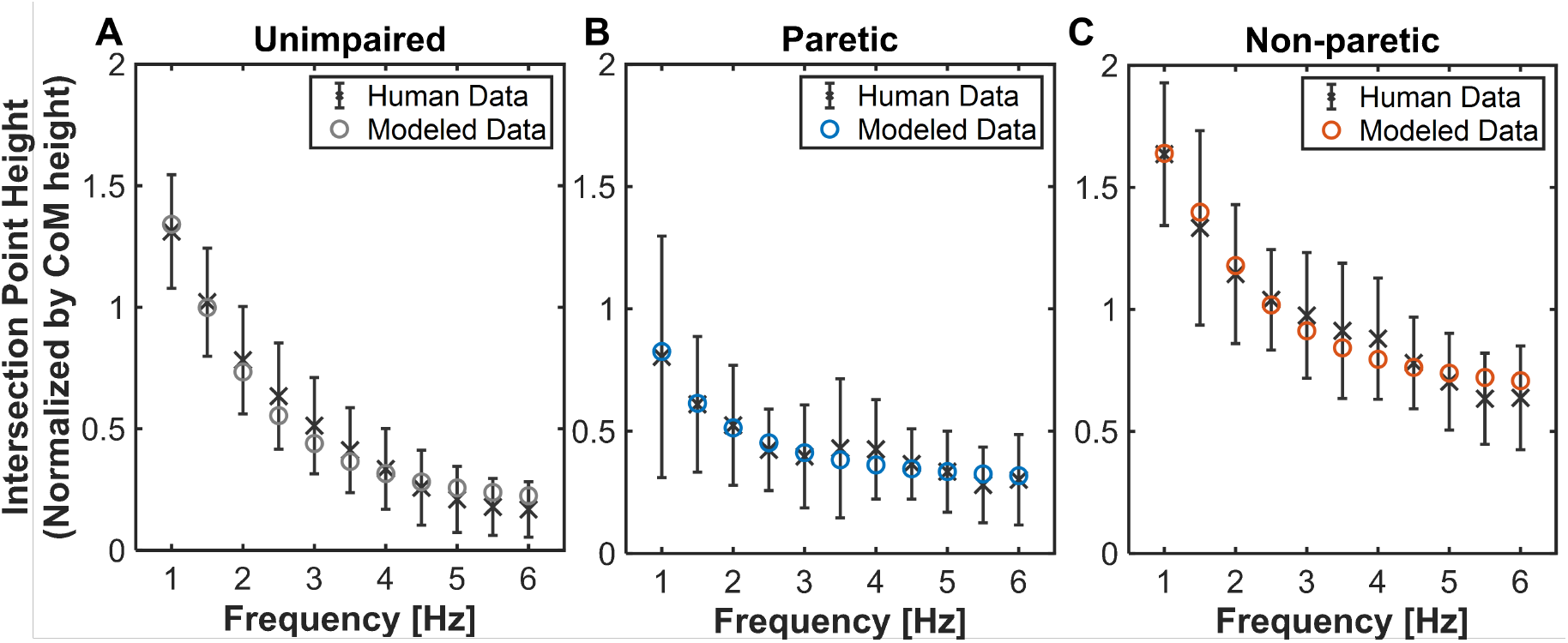
Normalized intersection point height of best-fit models (*⃝*) compared to experimental data (*×*; error bars indicate standard deviation) for A: unimpaired participants, B: paretic limb of post-stroke participants, C: non-paretic limb of post-stroke participants.

### 2.3 Optimal Gains

The best-fit control parameters determined the optimal controller gain matrix **K**_*LQR*_, composed of a 2 *×* 2 apparent stiffness matrix **K** and a 2 *×* 2 apparent damping matrix **B**:

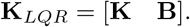

The optimal gain matrices corresponding to the best-fit models were:

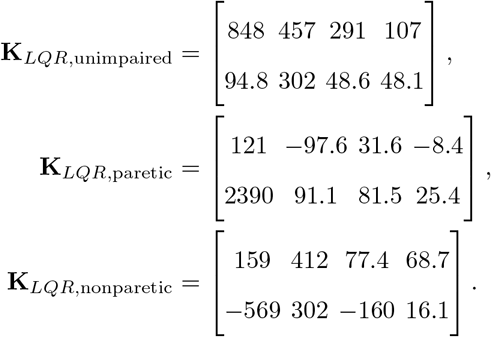

As in previous work [21], the apparent stiffness matrix was analyzed and compared between the three groups, as it provides insight on inter-joint coordination patterns. Specifically, the apparent stiffness matrix can be further decomposed into a symmetric component **K**_*S*_ and an antisymmetric component **K**_*A*_: while **K**_*S*_ has conservative properties and represents the biomechanical properties of the muscles as well as intra-joint and symmetric inter-joint neural feedback, a non-zero **K**_*A*_ may only arise from asymmetric inter-joint neural feedback [26, 27].

**K**_*S*_ and **K**_*A*_ identified for the three best-fit models were:

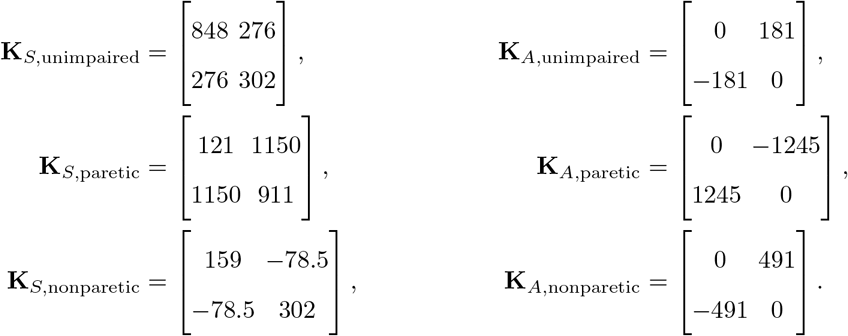

The ellipse-circle representation of 2-D stiffness allows visual comparison of the three apparent stiffness matrices (Figure 4) [21]: the ellipses represent **K**_*S*_, with the semi-major and semi-minor axes oriented along the principal and secondary eigenvectors in ankle-hip coordinates, respectively, and the axis lengths representing the corresponding eigenvalues; the circles represent **K**_*A*_, with the radius corresponding to the magnitude of its eigenvalues. The paretic-limb model had an antisymmetric component of stiffness that was much larger and in the opposite sense (negative hip-to-ankle coupling) compared to that of the unimpaired model (positive hip-to-ankle coupling), as well as a large symmetric component. The non-paretic limb showed a large antisymmetric component compared to the unimpaired model and in the same sense as that of the unimpaired model, while the symmetric component was much smaller than the other models.

**Fig. 4:**
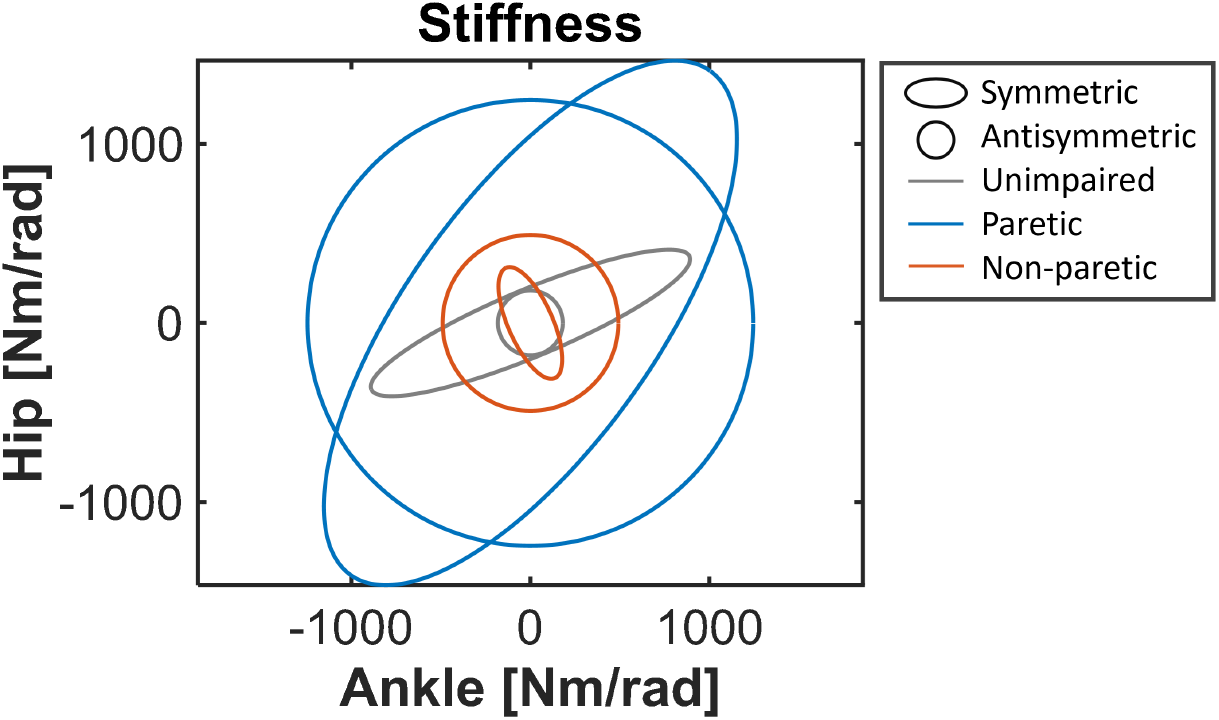
Symmetric component (ellipses) and antisymmetric component (circles) of the apparent stiffness matrix obtained from the best-fit models of unimpaired participants (gray), the paretic limb of post-stroke participants (blue), and the non-paretic limb of post-stroke participants (red).

The relative contribution of asymmetric inter-joint neural feedback can be quantified by the dimensionless ratio as defined by [28]:

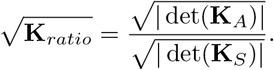

Note that this quantity does not depend on the choice of coordinates [21], and thus served as an appropriate measure to compare the extent to which asymmetric inter-joint neural feedback plays a role in the paretic versus non-paretic limbs of post-stroke participants and in the unimpaired older participants.

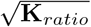 for each of the three best-fit models were:

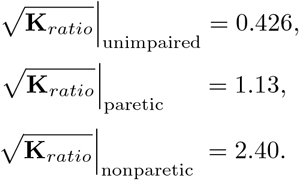

The non-paretic limb exhibited the largest 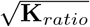, while the unimpaired model had the smallest, in agreement with the size of the circle relative to the ellipse in the graphical representation (Figure 4).

### 2.4 Distribution of Joint Torques

To compare inter-joint coordination patterns across the three conditions, the three best-fit models were simulated, and the distribution of commanded joint torques compared. Each model was simulated in MATLAB for 1000 seconds at 1000 Hz, to provide sufficient data points and reduce numerical errors due to discretization. The ankle and hip torques, *τ*_*a*_ and *τ*_*h*_, commanded at each time step were plotted in a 2D histogram in the (*τ*_*a*_, *τ*_*h*_) space (Figure 5). The torque distribution for the best-fit unimpaired-participants model exhibited a positive covariance between the ankle and hip torques (Figure 5A), suggesting positive coupling between the two joint torques. For the paretic-limb model, the torque distribution showed a negative covariance between the ankle and hip torques (Figure 5B), suggesting negative coupling. The non-paretic-limb model exhibited a positive covariance between joint torques (Figure 5C), as in the unimpaired model.

**Fig. 5:**
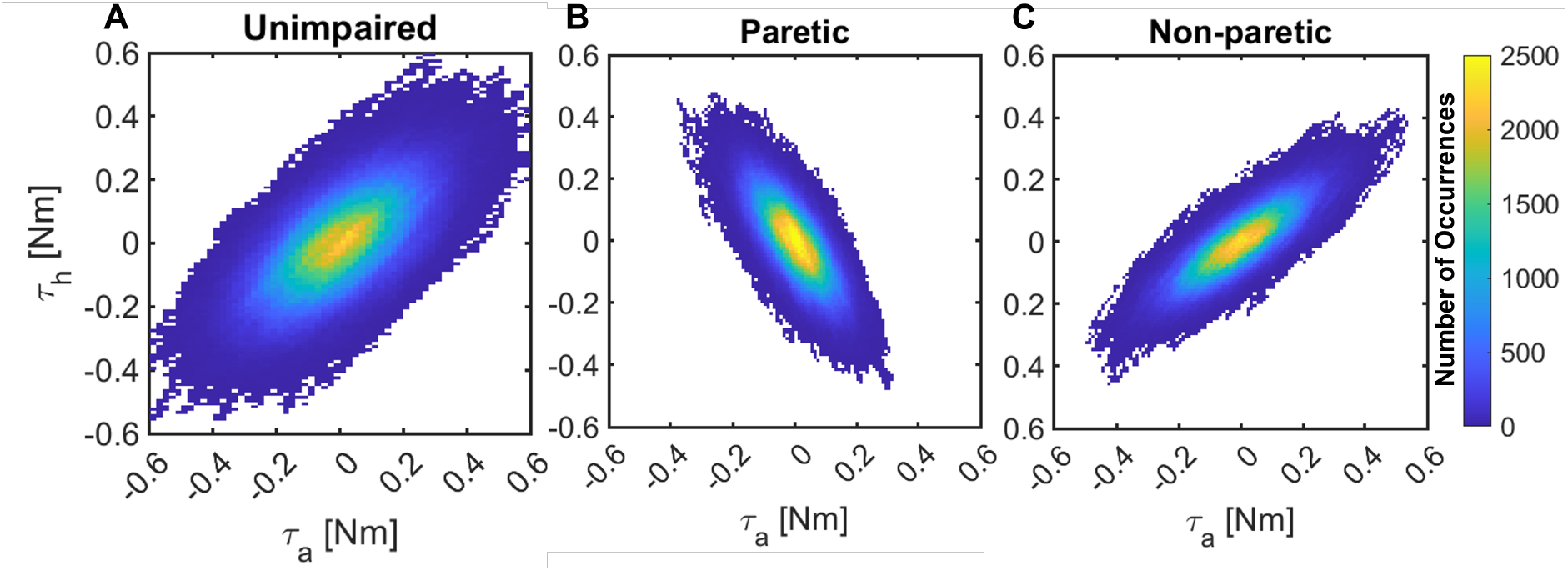
Distribution of commanded joint torques from simulations of best-fit models for A: unimpaired participants, B: the paretic limb of post-stroke participants, C: the non-paretic limb of post-stroke participants. The color bar shows the number of occurrences of the specific ankle and hip torque combination (*τ*_*a*_, *τ*_*h*_).

## 3 Discussion

Bartloff and colleagues observed notable differences in the intersection-point height between post-stroke individuals and similarly-aged controls [22]. Specifically, the non-paretic leg exhibited the highest overall intersection-point height across all frequencies, followed by the controls, and then the paretic leg. The researchers suggested that these differences may reflect disrupted coordination in the paretic limb and a compensatory strategy in the non-paretic limb. Motivated by these findings, in this study we modeled human balance control in an effort to identify apparent controllers that could describe the distinct patterns in human data.

A deliberately-simplified sagittal-plane quiet standing model stabilized by a linear full-state-feedback controller was able to account for the means of each leg of 12 individuals with history of stroke and of 22 unimpaired participants. To analyze balance control differences between the non-paretic and paretic limbs and between the post-stroke participant group and the control group, models were fit to the mean experimental intersection-point height data for each condition, and the best-fit models were compared. Results suggested an altered joint-torque coordination pattern and substantial asymmetric inter-joint neural feedback in the paretic leg compared to the controls. Furthermore, the non-paretic leg also exhibited substantially more reliance on neural feedback, possibly to compensate for the abnormal coordination pattern observed in the paretic limb (Figure 5B). These findings highlight the potential of the intersection point analysis as a quantitative assessment tool to evaluate balance control strategies that may lead to impairment.

In this study, the parameter *ω* was introduced to describe the degree of coupling between ankle and hip control. While the best-fit model describing the unimpaired group had a negative value of *ω*, suggesting positive coupling between the ankle and hip joint torques, the paretic-limb model had a positive *ω* value, suggesting negative coupling between the joint torques. This difference in joint coordination was also evident in the torque distribution plots (Figure 5), which showed a positive covariance for the unimpaired model and a negative covariance for the paretic-limb model. These results indicate that balance control in the paretic limb differed from the unimpaired participants, possibly due to compromised inter-joint coupling. Inter-joint coordination has previously been identified as a deficit in paretic leg coordination [16], and a causal relationship to behavioral deviations has been proposed [7].

This difference in balance control between the paretic-limb and unimpaired models was also evident when investigating the role of neural feedback in the apparent controller. To quantify neural contribution, the apparent stiffness matrices of the best-fit closed-loop systems were analyzed. The symmetric and antisymmetric components of the apparent stiffness matrix are related to distinct aspects of balance control: the symmetric component represents conservative properties, derivable from a potential function, whereas the antisymmetric component represents forces with curl, which are not derivable from a potential function [26, 27]. Physiologically, the symmetric component captures both intrinsic muscle stiffness and neural feedback; meanwhile, the antisymmetric component is non-zero only when asymmetric inter-joint neural feedback is present. In the current study, the antisymmetric component was non-zero for both the paretic and non-paretic limbs, as well as for the controls. This result is consistent with previous findings that intrinsic ankle muscle stiffness alone is insufficient to support standing balance [29, 30]. Additionally, prior research examining the role of asymmetric neural feedback in older and younger unimpaired adults also found that it was non-zero, further supporting asymmetric neural contributions to balance control [21].

When relative magnitudes of the antisymmetric and symmetric stiffness components were compared, the paretic limb exhibited a substantially larger ratio (1.13) compared to the controls (0.426). Furthermore, the antisymmetry was in the opposite sense, with the hip-to-ankle apparent stiffness being negative for the paretic-limb model, in contrast to the positive apparent stiffness in the controls. Unlike its conservative (spring-like) counterpart, the antisymmetric (rotational and non-spring-like) component of stiffness can, under certain conditions, act as an energy source and lead to instability. Therefore, there is a critical need for further research on the role of asymmetric neural feedback and its implications for balance control.

Previous studies provide evidence supporting these changes in balance control that occur in the paretic limb due to stroke. First, Varoqui and colleagues observed that post-stroke participants exhibited altered coordination patterns between the ankle and hip [31]. When tasked with reproducing ankle-hip coordination patterns that unimpaired individuals could produce with ease, the impaired participants either could not complete the task or exhibited significant instability when attempting to do so. Moreover, previous studies have also measured abnormal coupling across joints and abnormal muscle synergy patterns in the paretic leg [32–35]. For example, although evaluated in the supine position, maximum voluntary hip extension triggered involuntary ankle plantar-flexion [35]. This and related changes to joint coupling behavior in the paretic leg may be a result of altered heteronymous spinal pathways due to stroke [34]. These findings support the possibility that coupling between the ankle and hip joints was compromised in the post-stroke participants’ paretic limb.

Secondly, joint weakness may also contribute to altered coordination patterns. Hip joint torques for flexion and extension (sagittal plane), as well as knee and ankle joint torques, have been observed to be lower in the paretic limb compared to the non-paretic limb [36]. Furthermore, Twitchell reported that proximal joints tend to recover earlier than distal joints in post-stroke participants, which could lead to abnormal coupling between the hip and ankle joints [37].

Thirdly, evidence also suggests increased ankle stiffness in stroke participants. Studies using a custom ankle movement training device in chronic stroke participants (approximately 10 years post-stroke) reported a twofold increase in ankle plantar- and dorsi-flexion stiffness in the paretic limb compared to the non-paretic limb [38]. Meanwhile, other studies have observed increased passive ankle stiffness in dorsi-flexion [39, 40] but no significant changes in plantar-flexion [39]. Similarly, in the current study, relatively large values of apparent stiffness were observed in the paretic limb compared to the controls and the non-paretic limb. Some possible reasons for increased ankle stiffness are hypertonicity in the plantar-flexors [40] and altered intrinsic muscle properties induced by neurological damage [41]. Further investigation of the specific mechanisms underlying abnormal coordination patterns in the paretic leg is an important avenue for future work.

Shifting the focus to the non-paretic leg, a global parameter search of *ω* showed significant positive coupling between these joints, as was also apparent in the torque distribution plot. Notably, the non-paretic limb data could not be adequately fit without *ω*, underscoring the importance of coupling in the less affected leg. Nevertheless, the best-fit value of *ω* was relatively similar for both the non-paretic leg and unimpaired participants.

While best-fit *ω* and the corresponding torque distributions suggested a relatively similar degree of ankle-hip coupling in the non-paretic limb and controls, differences between the two conditions emerged when investigating the role of neural feedback in the apparent controller. The non-paretic limb exhibited a substantially larger ratio between the antisymmetric and symmetric components of the apparent stiffness (2.40) compared to the paretic limb (1.13) and the controls (0.426). This result suggested that a strong preference for asymmetric neural feedback was present in the non-paretic limb of post-stroke participants. Furthermore, the asymmetry was in the opposite sense compared to the paretic limb, with positive apparent stiffness in the hip-to-ankle direction, suggesting that this inclination may have emerged as a compensatory strategy, as proposed by Bartloff and colleagues [22].

It is possible that the post-stroke participants’ non-paretic leg compensated for the aforementioned coordination deficit observed in their paretic leg [42]. Supporting this idea, research on stroke-induced alterations in intermuscular coordination found that the paretic limb produced an anteriorly misdirected force when participants performed a seated lower extremity task (pushing on a pedal) [16]. While this task differed fundamentally from standing balance control, as it did not require maintaining whole-body postural stability and the CoP was constrained by the pedal pivot, such an abnormal force in the paretic limb would destabilize standing balance by generating a rearward-pitching angular acceleration [7]. This disruption necessitates a compensatory response that could involve increased reliance on neural feedback in the non-paretic limb.

Currently, post-stroke standing balance ability is primarily evaluated using clinical assessments such as the Fugl-Meyer Assessment of Lower Extremity [43], the Berg Balance Scale [44], and the Brunel Balance Assessment [45]. While these tests provide valuable insight into a patient’s functional ability, research has shown that functional balance ability and the presence of impairments do not correlate with each other after the acute phase of stroke [1, 10]. One reason is because patients are able to improve balance function through compensatory strategies, leaving impairments to persist ‘undercover’ [1, 4].

To assess impairments that affect balance function, clinicians commonly administer various tests. For example, weakness can be evaluated using the Motricity Index [46] and spasticity can be assessed with the Modified Ashworth Scale [47]. Additionally, prior studies that have examined balance impairment in stroke patients have typically used force plates to measure left-right-foot-force distribution and CoP motion [5, 48]. While these methods can partially assess balance impairment and can track rehabilitation progress, they do not distinguish whether improvements in standing balance stem from actual restoration of the paretic leg or compensatory strategies adopted by the non-paretic leg. A previous study that modeled responses to perturbations during balance found that the control effort exerted by the paretic leg was not simply a reflection of its weight-bearing contribution [14]. Similarly, Bartloff and colleagues showed that the observed differences in the intersection-point height between unimpaired and impaired individuals were not solely a function of left-right-foot-force asymmetry [22]. These findings suggest that foot-force asymmetry alone does not provide a sufficient characterization of balance impairment. Even when foot-force distribution is symmetrical, the non-paretic leg may still be compensating for deficits in the paretic leg. Thus, additional quantitative measures beyond foot-force magnitude distribution should also be considered to enhance therapy interventions. Specifically, deviations from a ‘normal’ intersection-point height pattern, as well as quantitative details on ankle-hip coordination and contributions of neural feedback, could serve as valuable indicators of impairment that could be addressed and restored through rehabilitation.

Employing a deliberately simplified model inevitably introduces limitations. The model used two degrees of freedom, representing the lower limbs and torso as rigid bodies. This structure was the simplest model that could reproduce the experimentally observed relationship between intersection-point height and frequency [18]. The model did not include the foot, and thus did not account for the height of the ankle above the ground. Although this results in an underestimate of the intersection-point height by the amount corresponding to the ankle height, analysis including an estimate of the ankle height showed that the main results were robust to the exclusion of the foot. The non-paretic and paretic legs of stroke participants were evaluated separately; thus, the coordination between the limbs could not be directly analyzed. In future studies, the non-paretic and paretic sides of the hips could be included in a single model for a more physiologically accurate description. Additionally, relative angles were selected as the coordinate frame for the ankle and hip joints. Care should be taken when interpreting the apparent stiffness and damping matrices, as they are dependent on this coordinate frame. However, the differences in the role of feedback were quantified using the ratio of the matrix determinants and do not depend on the coordinate frame.

Although the biomechanical model was nonlinear, the controller employed a linearized version of the model. As a result, the reported best-fit parameters are only valid for small motions around upright posture. This assumption is reasonable, as the experimental data consisted of participants standing quietly without large movements. We also assumed minimized control effort (large *α*) to reduce the search space of parameters. This choice was justified, as previous studies have shown that this assumption leads to better fit of human data [18, 19]. Furthermore, preliminary analysis showed that conclusions derived from comparing measures of interest were insensitive to the choice of *α*. Although this study focused on creating a simple descriptive model and controller, future work should explore other formulations that might also account for human data.

The model also used simplified neuromechanics. The stiffness reported in this study is the apparent stiffness, incorporating multiple factors such as intrinsic muscle mechanics and neural feedback. Moreover, the model does not explicitly account for delay, and all noise processes, including sensory noise, were lumped as additive motor noise. Despite these simplifications, the model was able to adequately account for the human data. Nevertheless, stroke can affect various aspects of sensorimotor control, and future studies should explore these components separately and in detail. As this effort could require more complex models with larger parameter spaces that incorporate factors such as sensory integration (proprioception, vision, and vestibular input), feedback delays, and muscle dynamics, the results of the current study could be used to simplify the search process of parameters that best fit human data. For example, the finding that a non-zero antisymmetric component best explains post-stroke human data could serve as a constraint to reduce the parameter search space.

Additionally, the sensitivity of the model’s parameters should be investigated further, particularly how these parameters change with varying intersection-point-height patterns observed across individuals. Although the findings of this study indicated altered coordination patterns in the paretic limb, the underlying impairments driving this behavior remain unclear. Investigating individual differences could reveal variations in ankle or hip dominance that contribute to abnormal joint coupling and lead to targeted therapeutic interventions. An extension of this work could explore how factors such as stroke onset time (acuteness), severity, and participant age affect the intersection-point height and thus control strategies. Validation studies are also needed to enhance the interpretability of the best-fit parameters.

Mathematical modeling goes beyond accounting for differences from a baseline; it provides quantitative details of the underlying control strategies driving those differences. In this study, the height of the intersection point of foot-ground force lines of action and its relation to frequency were simulated using a simple, interpretable model to compare quiet balance control strategies across post-stroke individuals and similarly-aged controls. Results revealed that ankle-hip torque coordination was altered in the paretic limb compared to similarly-aged controls. Furthermore, our findings suggest that the non-paretic limb may compensate for impairments in the paretic limb through increased neural feedback. As one of the first studies to model quiet balance control post-stroke, this work lays the foundation for future modeling efforts and paves the way for a deeper understanding of balance impairments caused by stroke.

## 4 Methods

### 4.1 Experimental Data

The experimental protocol is summarized briefly here, as complete details are available in the published studies which provided the data used in this study [22, 24].

The post-stroke data from [22] included 12 participants (4 female, age: 62*±*14 years, mean *±* standard deviation) over 6 months post-stroke with chronic hemiplegia who could stand unassisted for at least five minutes with no pain. The public data set [24] included data from both younger and older adults standing under various conditions (eyes open and closed, standing on foam versus rigid floor). The data from 22 unimpaired older participants (11 female, age: 69 *±* 11 years) with eyes open and standing on a rigid surface were used as the control data for comparison.

Sagittal-plane foot-ground force data (center-of-pressure location and force vector orientation) were collected at 100 Hz during the quiet standing trials, in which participants stood with each foot on a force plate, arms by their sides, looking at a target straight ahead [22, 24]. The post-stroke data consisted of one 15-s trial for nine of the participants, and one 60-s trial for three of the participants, while the control data consisted of three 60-s trials [22, 24]. Preliminary analysis conducted by Bartloff and colleagues showed that the intersection point computed from data segments of 10 s or more showed the same pattern, thus justifying the analysis with varying data length [22].

#### 4.1.1 Intersection-Point Analysis

In the present study, the intersection-point height was computed using a revised method that ensures the measure obtained is scale invariant [25]. Furthermore, in contrast to [22] which expressed the center of pressure (CoP) relative to the center of mass (CoM), in this study CoP was expressed relative to the lab frame to allow for comparison with previous analyses using the same approach [18, 19, 21].

The sagittal-plane intersection-point height was computed from the force data of each limb of each participant, for each trial, at 11 distinct frequencies from 1 Hz to 6 Hz at 0.5-Hz increments; 1 Hz was chosen as the lowest frequency bin, as the intersection-point height computed at lower frequencies showed high inter-subject variance. The procedure can be summarized as follows:

1. the time series data of foot-ground force orientation (*θ*_*F*_) and CoP displacement (*x*_*CoP*_) were band-pass filtered with a 2nd-order zero-lag Butterworth filter with a bandwidth of 0.5 Hz, centered at the frequency of interest (each of the 11 frequencies); the first and last 6% of the band-pass-filtered time-series data were ignored to remove transients from the filtering;
2. the intersection-point height was computed as the covariance between the band-pass-filtered *θ*_*F*_ and *x*_*CoP*_ divided by the variance of the band-pass-filtered *θ*_*F*_;
3. the intersection-point height was then normalized by the CoM height (estimated as 56% of the total height [49]);
4. steps 1 to 3 were repeated for each of the 11 frequencies to produce the intersection-point-height curve with respect to frequency.

Average intersection-point-height curves were then obtained for the paretic limb of post-stroke participants, the non-paretic limb of post-stroke participants, and the unimpaired control participants by averaging all the intersection-point-height data at each frequency from all the trials of all subjects in the respective groups. For the unimpaired control participants, the intersection-point-height curve computed from the dominant-leg data and non-dominant-leg data showed similar trends [22] and therefore were averaged to produce a single intersection-point-height curve for the control group. Any data points that were three standard-deviations away from the group mean were removed, and the mean intersection-point height re-computed.

### 4.2 Modeling

The modeling approach used in this work followed a similar procedure to that of previous studies [18, 19, 21] with some modifications to the controller parameters to enable satisfactory fit to the intersection-point curve of the non-paretic limb.

#### 4.2.1 Biomechanical Model

In this work, as in the previous studies, the simplest descriptive model that could reproduce the experimental intersection-point data was used to obtain the apparent control strategies employed in the impaired versus unimpaired limbs.

Given that a single-inverted-pendulum model of standing was unable to reproduce the universally-observed frequency-varying intersection-point height, with heights below the CoM height at higher frequencies [18], a double-inverted-pendulum model was employed. The ankle and hip joint angles (*q*_*a*_, *q*_*h*_) and angular velocities 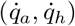 were chosen as the state variables 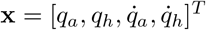, and the ankle and hip joint torques as the input *τ* = [*τ*_*a*_, *τ*_*h*_]^*T*^ (Figure 2).

Table 1 shows the lumped model parameters which were determined based on the average mass and height of the post-stroke participants (86.0 kg, 1.69 m) and those of the unimpaired participants in the experimental studies (68.7 kg, 1.61 m), along with the anthropometric distribution of young male adults from [50]. The mass and height of the feet were neglected, as the feet remained stationary. The CoM position for the lower body (link 1) is given with respect to the ankle joint, while that for the upper body (link 2) is given with respect to the hip joint. The moment of inertia is given with respect to the CoM of each link. The equations of motion, center-of-pressure position, and foot-ground force vector orientation were computed as in previous work [18].

The controller-specified ankle and hip joint torques were modulated by white, mutually uncorrelated, zero-mean normally distributed noise processes, with standard deviations *σ*_*ankle*_ and *σ*_*hip*_, respectively. These noise processes accounted for the lumped stochasticity in the system, including the effects of sensory, motor, and computational noise, as perceived at the level of joint torques. The relative strength of the two apparent motor noise processes was described by the noise ratio *σ*_*r*_ = *σ*_*ankle*_*/σ*_*hip*_, and was one of the free parameters that were varied to fit the human intersection-point-height data.

#### Controller

Following the approach in the previous studies [18, 19, 21], the linear quadratic regulator (LQR) was chosen to describe the apparent controller. The double-inverted-pendulum equations of motion were linearized about the upright pose (**x**_0_ = [0, 0, 0, 0]^*T*^, *τ*_0_ = [0, 0]^*T*^) and the LQR control gain matrix **K**_*LQR*_ *∈* ℝ^2*×*4^ was computed to minimize the cost (*J*) given by:

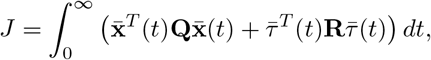

where 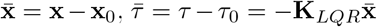, **Q** *∈* ℝ^4*×*4^ is the state-cost weighting matrix, and **R** *∈* ℝ^2*×*2^ is the control-input-cost weighting matrix. Thus, the state-cost and control-input-cost weighting matrices, **Q** and **R**, determined the controller gains.

In previous work [18, 19, 21] the control-input-cost weighting matrix **R** was parametrized as a diagonal matrix:

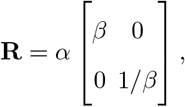

where *α >* 0 determined the weight on the input cost relative to the state cost, and *β >* 0 determined the weight on the ankle-torque cost relative to the hip-torque cost. However, preliminary analysis revealed that the intersection-point-height data of the non-paretic limb could not be described by a controller restricted to a diagonal input-cost weighting matrix. Thus, in this work, the off-diagonal elements of **R** were introduced as an additional parameter:

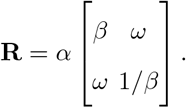

*ω* influences the coordination between the ankle and hip torques: *ω <* 0 encourages positive coupling between the ankle and hip torques, while *ω >* 0 encourages negative coupling. *ω* was restricted to |*ω*| *<* 1 to ensure that **R** was a positive definite matrix, as required by the LQR procedure.

To find the simplest competent descriptive model with the smallest number of parameters, the state-cost weighting matrix **Q** was fixed to be the identity matrix as in previous studies [18, 19, 21]. This 7 simplification is justified as long as the input cost is weighted much more than the state cost (i.e., 8 9 |**R**| *≫* |**Q**|), such that the contribution of the state cost (and thus **Q**) is negligible; |**R**| *≫* |**Q**| can be achieved by choosing a sufficiently large *α*. A sensitivity analysis on the parameter *α* showed that the conclusions drawn from comparing the best-fit controllers for the different conditions (unimpaired, paretic, non-paretic) using the key measures of interest (see 4.3) were robust to changes in *α*. Thus, *α* was fixed as a large constant value (*α* = 10^6^), leaving *β* and *ω* among the free parameters varied to fit the experimental intersection-point-height curves. Whether *α* = 10^6^ was large enough was evaluated by checking the closed-loop poles of the system: if their location in the complex plane corresponded to one in which the open-loop unstable poles were reflected across the imaginary axis, then the condition |**R**| *≫* |**Q**| was met. For all three cases (unimpaired, paretic, non-paretic), the best-fit parameters yielded closed-loop poles that were a reflection of the open-loop unstable poles across the imaginary axis, thus meeting the condition that the input cost dominated the cost function.

#### 4.2.3 Best-Fit Control Parameter Search

The goal of this work was to quantify and compare the apparent control strategies employed by the post-stroke and similarly-aged unimpaired participants. To this end, a global parameter search was conducted over the controller parameters *β* and *ω*, and the apparent-motor-noise ratio *σ*_*r*_, to determine which parameter combination best reproduced experimental intersection-point-height curves of the paretic and non-paretic limbs of post-stroke participants, and that of unimpaired participants.

For each of the three experimental intersection-point-height curves (unimpaired, paretic, non-paretic), the following sets of parameters were tested: 1

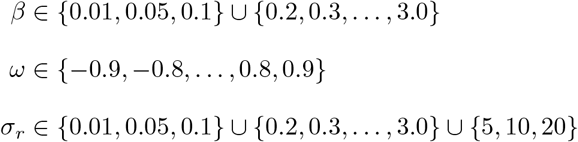

Note that *β* and *σ*_*r*_ were restricted to positive values, while *ω* was restricted to be within the interval 1 (*−*1, 1) as required for the LQR cost function.

For each combination of parameters, the intersection-point-height curve was estimated using an analytical method based on the linearized double-inverted-pendulum model [23, 25]. The intersection-point height at each frequency was normalized by the CoM height (56% of the average height for each subject group) to maintain consistency with experimental methods.

The goodness-of-fit of the model’s intersection-point-height curve was quantified by the root-mean-square error (RMSE) between the model curve and the average experimental curve of the given participant group and limb under consideration:

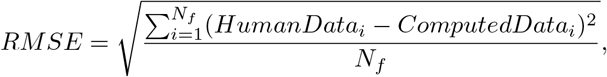

where *HumanData*_*i*_ was the average intersection-point height of the human data (for the paretic limb, non-paretic limb, or unimpaired participants) at the *i*th frequency band, *ComputedData*_*i*_ was the intersection-point height computed from the model at the *i*th frequency band, and *N*_*f*_ = 12 was the total number of frequency bands. The best-fit model corresponded to the model with the lowest *RMSE* value.

### 4.3 Model Comparisons

The best-fit models for the post-stroke paretic limb, post-stroke non-paretic limb, and unimpaired participants were compared to identify differences in the apparent control strategies between the different groups and limbs. Note that there is a complex interplay between the parameters *β* and *ω* in the input cost. Thus, care should be taken when interpreting results, and the best-fit parameters alone may not fully explain the differences between the populations and legs. To gain a holistic view of coordination patterns, the output controller gains and the resulting closed-loop dynamics were analyzed using measures outlined in this section.

#### 4.3.1 Best-Fit Controller Gains

The apparent controller for each of the best-fit models was represented by the LQR gain matrix **K**_*LQR*_. The gain matrix was composed of two 2 *×* 2 matrices that can be interpreted as the apparent stiffness matrix **K** and the apparent damping matrix **B**:

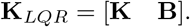

**K**, which is the gradient of the joint torque vector field, can be further decomposed into symmetric and 6 antisymmetric components, **K**_*S*_ and **K**_*A*_:

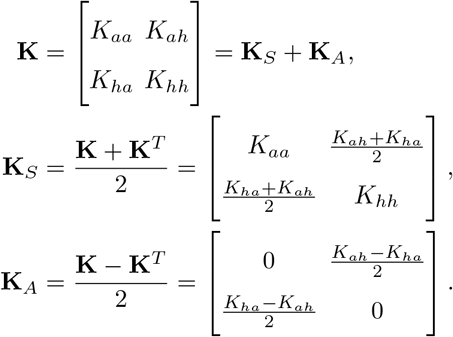

This decomposition of the apparent stiffness into its symmetric and antisymmetric components is insight-3 ful because **K**_*S*_ and **K**_*A*_ capture the contributions of different feedback mechanisms: **K**_*S*_ describes the net effect of intrinsic muscle impedance, intra-joint neural feedback, and symmetric inter-joint neural feedback, while **K**_*A*_ represents contributions from asymmetric inter-joint neural feedback [26, 27]. This can be explained by the fact that, under constant neural input, the net muscle force can be written as the gradient of the stored elastic energy, i.e. a scalar potential function; thus, if intrinsic muscle stiffness alone is responsible for producing joint torques, the curl of the torque vector field (which is the gradient of a scalar potential function in this case) must be zero, implying that **K**_*A*_ = 0. Intra-joint and symmetric inter-joint neural feedback also result in a torque vector field with zero curl, contributing to symmetric stiffness. A non-zero antisymmetric component of stiffness can only arise from joint torques resulting from asymmetric inter-joint neural feedback.

Note that, the symmetric and antisymmetric components of the apparent damping matrix cannot be 1 attributed to different feedback mechanisms in a similar way [21]. Therefore, the analysis presented here focused on comparing the relative size of the stiffness components.

The contributions of **K**_*S*_ and **K**_*A*_ were represented graphically in ankle-hip coordinates using the ellipse-circle representation as in [21]. The symmetric matrix **K**_*S*_ was represented by an ellipse with its semi-major axis oriented along its principal eigenvector in ankle-hip coordinates, and length determined by the principal eigenvalue, while the semi-minor axis was oriented along the second eigenvector with 1 length determined by the corresponding eigenvalue. The antisymmetrix matrix **K**_*A*_ was represented by a circle with radius equal to the magnitude of the off-diagonal elements (which correspond to the magnitude of its eigenvalues). This visualization allowed for qualitative comparison of the relative size of each stiffness component.

The relative contribution of asymmetric inter-joint neural feedback in each model was also quantitatively assessed by computing a dimensionless ratio between the magnitude of **K**_*A*_ and that of **K**_*S*_, as defined by [28]:

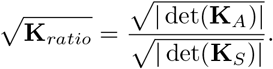

#### 4.3.2 Distribution of Joint Torques

The closed-loop dynamics of the best-fit model for each group (unimpaired, paretic, non-paretic) was simulated to analyze the relative contribution of the ankle and hip torques and the coordination between the two joints. The models were simulated in MATLAB at 1000 Hz using semi-implicit Euler integration, for one trial of 1000 seconds initialized at **x**_0_. A high simulation frequency of 1000 Hz was chosen to reduce the discretization error. The motor noise levels were normalized such that

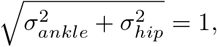

to produce comparable postural sway for all three model simulations. The commanded ankle and hip torques, *τ*_*a*_ and *τ*_*h*_, were visualized as 2D histograms in the (*τ*_*a*_, *τ*_*h*_) space to assess the coordination patterns between the two joints and how they differed across the three best-fit models.

## Data Availability

All data supporting the findings of this study are available within the paper. Additional datasets, including processed foot-force signals and model outputs, are available to reviewers and will be made publicly available in a GitHub repository upon publication. Portions of the dataset were adapted from a previously published study [22] and a publicly available dataset [24].

## Code availability

All code used for data analysis, modeling, and figure generation is available to reviewers and will be made publicly available in a GitHub repository upon publication. The repository includes preprocessing routines, model-fitting algorithms, and plotting scripts to reproduce the results presented in this study.

## Acknowledgments

This study was funded in part by the 2023–2024 MathWorks Mechanical Engineering Fellowship (to K.S.); National Science and Engineering Research Council of Canada Postgraduate Scholarships—Doctoral (to R.S.-D.); V. Horne Henry Fund, University of Wisconsin (to K.G.G.); Marsh Fund, University of Wisconsin (to K.G.G.); Eric P. and Evelyn E. Newman Fund (to N.H.); and Gloria Blake Endowment Fund (to N.H.);.

## Author Information

### Contributions

K.G.G. and N.H. conceived and designed research; K.S. and R.S.-D. performed modeling and simulation; K.S., R.S.-D., and K.G.G. analyzed data; K.S., R.S.-D., K.G.G., and N.H. interpreted results; K.S. and R.S.-D. prepared figures; K.S. and R.S.-D. drafted manuscript; K.S., R.S.-D., K.G.G., and N.H. edited 1 and revised manuscript; K.S., R.S.-D., K.G.G., and N.H. approved final version of manuscript.

